# The RNA Binding Protein FMRP Promotes Myelin Sheath Growth

**DOI:** 10.1101/669895

**Authors:** Caleb A. Doll, Katie M. Yergert, Bruce H. Appel

## Abstract

During development, oligodendrocytes in the central nervous system extend a multitude of processes that wrap axons with myelin. The highly polarized oligodendrocytes generate myelin sheaths on many different axons, which are far removed from the cell body. Neurons use RNA binding proteins to transport, stabilize, and locally translate mRNA in distal domains of neurons. Local synthesis of synaptic proteins during neurodevelopment facilitates the rapid structural and functional changes underlying neural plasticity and avoids extensive protein transport. We hypothesize that RNA binding proteins also regulate local mRNA regulation in oligodendrocytes to promote myelin sheath growth. Fragile X mental retardation protein (FMRP), an RNA binding protein that plays essential roles in the growth and maturation of neurons, is also expressed in oligodendrocytes. To determine whether oligodendrocytes require FMRP for myelin sheath development, we examined *fmr1^-/-^* mutant zebrafish and drove *FMR1* expression specifically in oligodendrocytes. We found oligodendrocytes in *fmr1^-/-^*mutants developed myelin sheaths of diminished length, a phenotype that can be autonomously rescued in oligodendrocytes with *FMR1* expression. Myelin basic protein (Mbp), an essential myelin protein, was reduced in myelin tracts of *fmr1^-/-^* mutants, but loss of FMRP function did not impact the localization of *mbpa* transcript in myelin. Finally, expression of FMR1-I304N, a missense allele that abrogates FMRP association with ribosomes, failed to rescue *fmr1^-/-^* mutant sheath growth and induced short myelin sheaths in oligodendrocytes of wild-type larvae. Taken together, these data suggest that FMRP promotes sheath growth through local regulation of translation.

## Introduction

Oligodendroglia in the central nervous system simultaneously wrap numerous axons in a highly dynamic ensheathment process that takes place at a distance from the nucleus [1]. Mature oligodendrocytes can ultimately form more than a dozen stabilized myelin sheaths per cell, with variability in the lengths of individual myelin internodes [2, 3]. How can these complex cells coordinate so many independent axonal interactions from a distance? Subcellular localization of mRNA represents a powerful mechanism to achieve on demand, localized protein production [4]. In the nervous system, this phenomenon allows cells to achieve remarkable plasticity, despite physical separation from the cell body [5, 6]. Oligodendrocytes express an array of RNA binding proteins (RBPs) [7–10], which may serve to regulate mRNA at active sites of ensheathment to meet independent protein requirements in each myelin sheath.

Fragile X syndrome (FXS), the most common heritable form of autism spectrum disorder [11], is caused by transcriptional silencing of the *FMR1* gene, which encodes the RBP Fragile X mental retardation protein (FMRP) [12]. FXS can be comorbid with epilepsy, ADHD, and anxiety [13–16], and FMRP plays prominent roles in a wide range of neural functions, including neural plasticity [17] and synaptic development [18–20]. Importantly, longitudinal neuroimaging has documented region-specific white matter abnormalities in individuals with FXS [21, 22], thereby also implicating oligodendrocytes in the disease state.

We hypothesize that FMRP regulates mRNA in oligodendrocytes to facilitate myelin sheath growth. As an RBP, FMRP has been implicated in mRNA stabilization [23], transport [24], and activity-dependent translation [25, 26] of critical synaptic genes in neurons; functions that could also facilitate local protein synthesis in highly polarized oligodendrocytes. Glial cells express FMRP during early brain development [9, 27], and *Fmr1* knockout mice show delayed cortical myelination during early postnatal stages [28]. However, the relative glial contribution of FMRP in myelination remains unclear. We used zebrafish to explore roles for FMRP in the early developmental myelination of axons, utilizing cell type-specific methods to visualize and manipulate *fmr1* and associated target genes in vivo, finding crucial FMRP requirements in myelin sheath growth and regulation of myelin genes.

## Results

### FMRP is localized within nascent myelin sheaths

Although the bulk of FMRP studies have focused on neuronal development and function, oligodendrocyte lineage cells also express FMRP [9, 27]. Our own RNA-Seq study revealed that zebrafish oligodendrocyte precursor cells (OPCs) and myelinating oligodendrocytes express *fmr1* (Fig. 1A; [29]) and RNA-Seq analysis of cells isolated from mouse brain indicated that OPCs and newly formed oligodendrocytes express *Fmr1* at levels comparable to neurons (Fig. 1B; [30]). In order to achieve distal regulation of target mRNAs, FMRP must be localized to myelin sheaths. To determine the subcellular localization of FMRP in live zebrafish larvae, we used Tol2-based transgenesis to transiently express a human FMR1-EGFP fusion protein behind *sox10* regulatory elements, which drive expression in OPCs and oligodendrocytes [31]. We injected *sox10:*FMR1-EGFP into 1-cell transgenic *sox10:*mRFP embryos expressing the oligodendrocyte membrane reporter *sox10:*mRFP [32] and subsequently examined individual cells at 4 days post-fertilization (dpf). In oligodendrocytes of the dorsal and ventral spinal cord, we noted bright FMR1-EGFP expression in cell bodies and more punctate expression in myelin sheaths (1C-D), including the terminal ends of RFP+ myelin sheaths (Fig. 1C’,D’, inset). These data demonstrate subcellular localization of FMRP in myelin sheaths during early developmental myelination.

**Figure 1.**
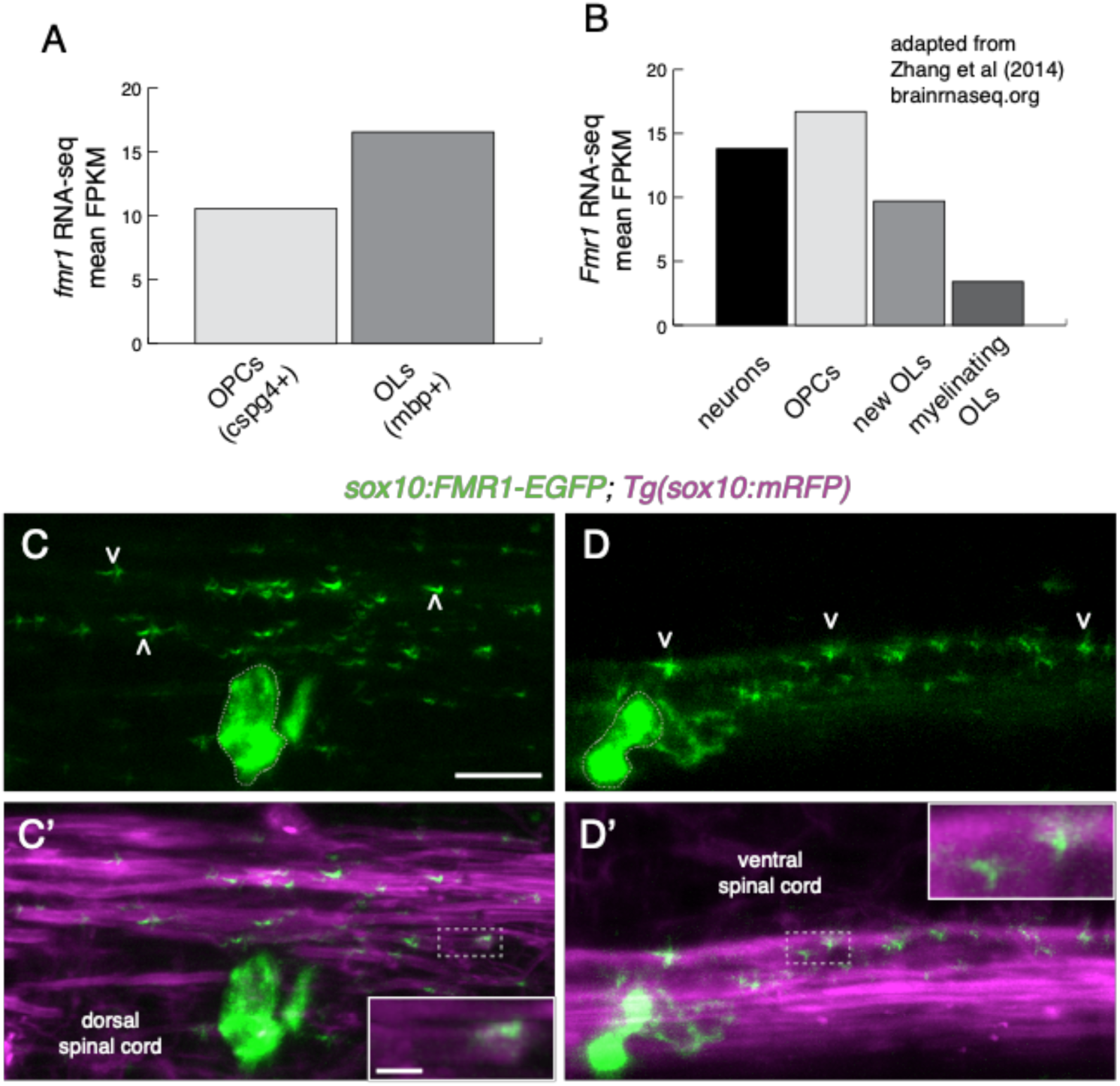
FMRP is localized within nascent myelin sheaths. (**A**) Zebrafish *fmr1* expression levels (fragments per kilobase of transcript per million mapped reads; FPKM) from RNA-Seq of FAC-sorted *cspg4*+*olig2*+ OPCs and *mbpa*+*olig2*+ oligodendrocytes [29]. (**B**) Murine *Fmr1* expression (FPKM) from RNA-Seq of neurons and oligodendrocyte lineage cells (adapted from [30]; brainseq.org). (**C-C’**) Lateral images of a living 4 dpf *Tg(sox10:mRFP)* transgenic larva transiently expressing *sox10:*FMR1-EGFP, a human FMR1-EGFP fusion construct, in an oligodendrocyte in the dorsal spinal cord. FMR1-EGFP expression is highest in the cell body (dashed outline), with dimmer, punctate expression noted in *sox10:*mRFP+ myelin sheaths of the dorsal spinal cord (arrowheads), including the terminal ends of sheaths (**C’**, inset). (**D,D’**) An oligodendrocyte in the ventral spinal cord transiently expressing FMR1-EGFP in a single myelin sheath on a large diameter Mauthner axon. The fusion protein is widespread in the cell soma (dashed outline) and punctate throughout the myelin sheath (arrowheads). FMR1-EGFP co-localizes with the *sox10:*mRFP+ myelin membrane (**D’** inset). Wide scale bar=10 µm, inset scale bar=2 µm.

### Global loss of FMRP leads to reduced myelin sheath growth and dynamics

FMRP-deficient neurons display prominent abnormalities in synaptic structure, with immature dendritic spines representing the hallmark phenotype in FXS patients [33]. Does FMRP also promote the maturation of myelin sheaths during development? With evidence of subcellular localization of FMRP within developing myelin sheaths, we next examined requirements for FMRP in oligodendrocyte development. Larvae homozygous for the loss-of-function *fmr1^hu2787^* allele do not display any obvious physical defects, and grow into viable, fertile adults [34]. To examine individual oligodendrocytes in the developing spinal cord, we mosaically expressed an oligodendrocyte-specific membrane tethered reporter, *mbpa*:EGFP-CAAX, which distinctly labels myelin sheaths [35]. We injected *mbpa:*EGFP-CAAX into 1-cell wild-type and maternal-zygotic *fmr1^hu278^* (*mzfmr1^-/-^*) embryos, in which any potential maternal contribution of FMRP is absent. We then quantified the length of individual myelin sheaths at 4 dpf, finding that sheaths in *mzfmr1^-/-^* loss-of-function larvae were reduced ∼35% in length compared to wild-type (Fig. 2C). Oligodendrocytes in *mzfmr1^-/-^* larvae formed approximately the same total number of sheaths per cell as wild-type (Fig. 2D). Taken together, the cumulative sheath length per cell was dramatically reduced in the absence of FMRP (Fig. 2E). With prominent deficits in overall myelin sheath growth at 4 dpf, these data suggest that FMRP promotes the growth of myelin sheaths.

**Figure 2.**
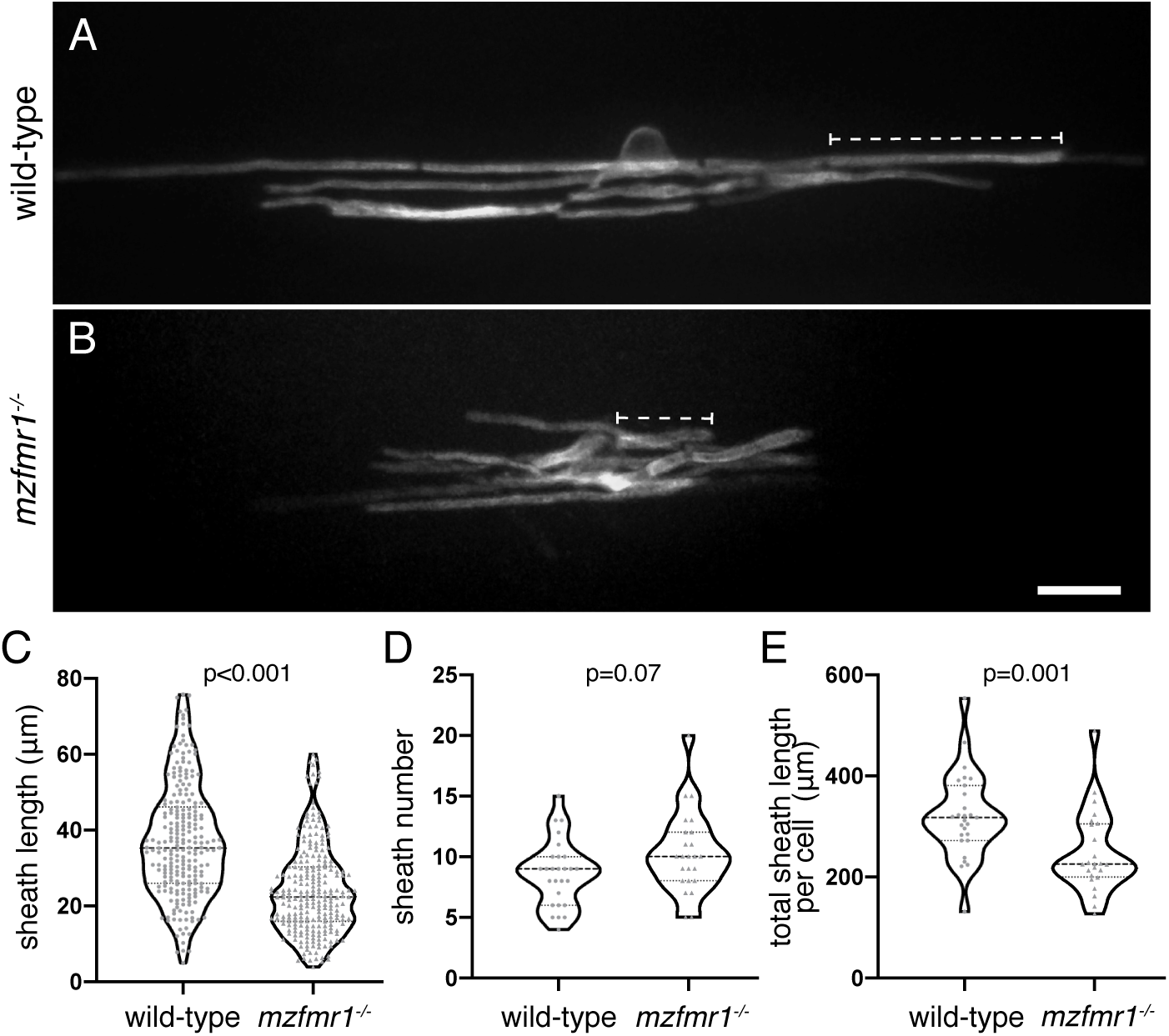
Global loss of FMRP leads to reduced myelin sheath length. Lateral images of oligodendrocytes in the spinal cord of living wild-type (**A**) and *mzfmr1^-/-^* (**B**) larvae labeled by *mbp:*EGFP-CAAX. The *mbp:*EGFP-CAAX reporter distinctly labels individual myelin sheaths (brackets). Average sheath length (**C**), sheaths per cell (**D**), and total cumulative sheath length (**E**) in wild-type and *mzfmr1^-/-^*loss-of-function mutants. Violin plot centerlines represent median values and hinges represent 25^th^ and 75^th^ percentiles, with all data points shown. Wild-type: n=234 sheaths, 27 cells; *mzfmr1^-/-^*: n=247 sheaths, 24 cells. Significance determined by Mann-Whitney tests. Scale bar=10 µm.

The short myelin sheaths noted in *fmr1^-/-^*larvae could result from deficient growth or from sheath instability and excess retraction. We examined these possibilities by capturing time-lapse images of the developing spinal cord at 3 dpf, when sheath development is highly dynamic in newly differentiated oligodendrocytes (Fig. 3). We recorded three-dimensional images of transgenic wild-type and *fmr1^-/-^* loss-of-function larvae expressing *sox10:*mRFP every 15 minutes for 4 hours. We then quantified the length of individual myelin sheaths at each time point and analyzed the overall growth or retraction (t240 - t0) as well as the dynamic range (maximum value – minimum value), the latter a measure of overall capacity for growth or retraction. During this 4-hour imaging window, we found that wild-type myelin sheaths gained an average of 1.34±0.23µm in length, whereas *fmr1^-/-^* mutant sheaths displayed very little positive growth (0.33±0.22µm, Fig. 3B). In addition, wild-type sheaths were highly dynamic, with an overall average range of 2.47±0.2µm. In contrast, *fmr1^-/-^* sheaths displayed an average range of 1.9±0.14µm, a 23% reduction compared to wild-type (Fig. 3C). In summary, these data indicate an oligodendrocyte requirement for FMRP in the highly dynamic phase of initial myelin sheath growth, with reduced sheath growth and dynamic capacity in *fmr1^-/-^* mutants.

**Figure 3.**
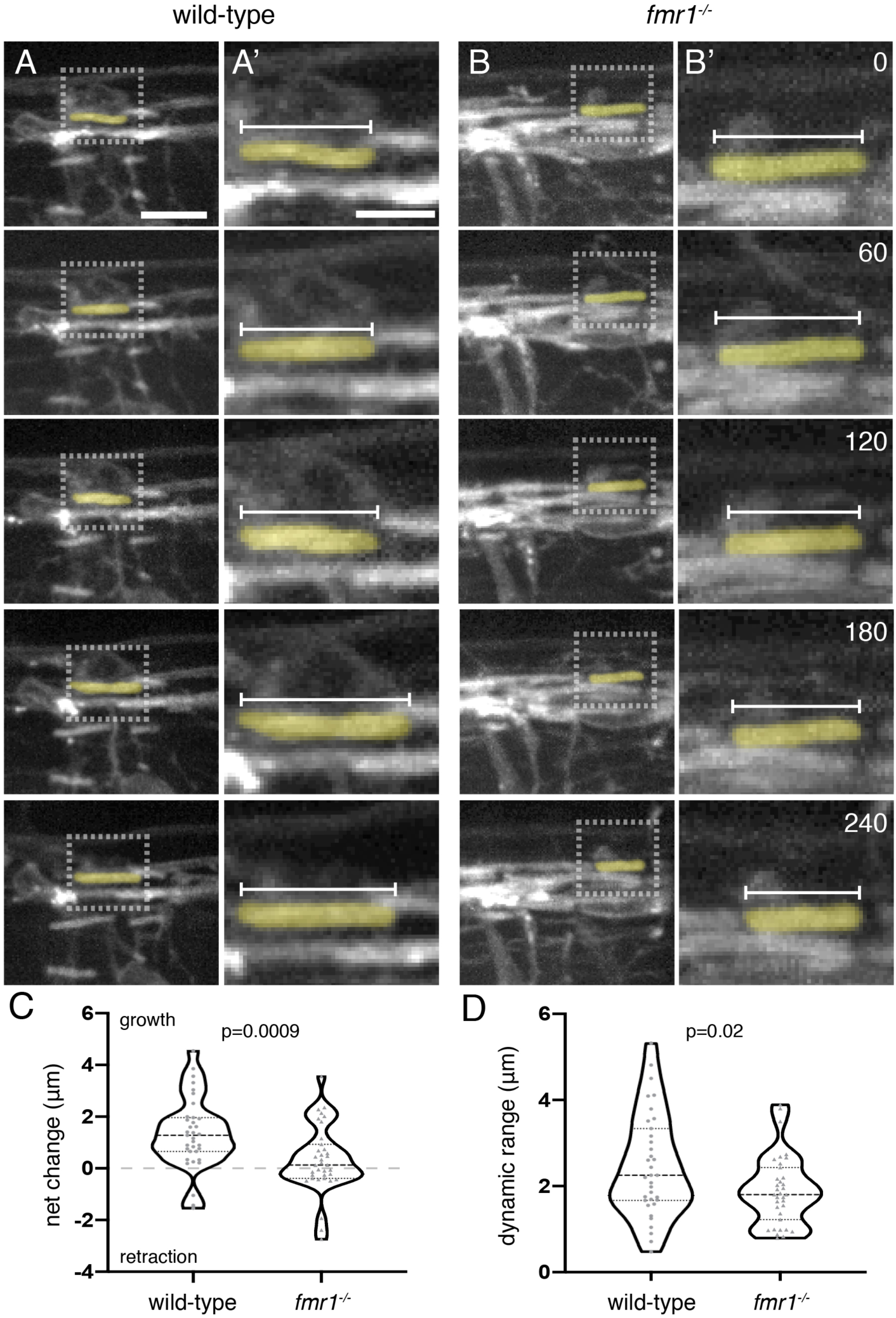
FMRP promotes dynamic myelin sheath growth. *sox10*:mRFP+ myelin sheaths in the dorsal spinal cord of transgenic *Tg(sox10:mRFP)* wild-type (**A**) and *fmr1^-/-^* mutant (**B**) larvae at 3 dpf. Lateral three-dimensional images of the spinal cord were captured every 15 minutes for 4 hours (each row represents an hourly time point; scale bar=10 µm). (**A’**,**B’**) High magnification detail of individual *sox10*:mRFP+ myelin sheaths, inset in **A** and **B** (scale bar=5 µm). Numbers in top right corners denote time elapsed from initial time point. (**C**) Change in average length of individual myelin sheaths (t240-t0; wild-type=35 sheaths, 8 larvae; *fmr1^-/-^*=35 sheaths, 7 larvae; Mann-Whitney test). Positive values indicate sheath growth and negative values denote retraction from the initial timepoint. (**D**) Dynamic range of individual myelin sheaths (maximum value-minimum value) Statistical significance evaluated using unpaired *t* test.

### FMRP autonomously promotes myelin sheath growth

Because *fmr1^hu2787^* mutants globally lack FMRP, deficient sheath growth could stem from dysfunction in neurons or oligodendrocytes. To test whether FMRP functions autonomously to promote myelin sheath growth, we transiently expressed human *FMR1* using *myrf* regulatory DNA, which drives expression in differentiated oligodendrocytes [36]. In contrast to the fusion protein encoded by *sox10:*FMR1-EGFP presented in Fig. 1, FMR1 and EGFP-CAAX are produced as distinct peptides from the *myrf:*FMR1-IRES-EGFP-CAAX construct. We injected control *myrf:*EGFP-CAAX and FMR1 plasmids in both wild-type and *mzfmr1^-/-^*loss-of-function embryos at the 1-cell stage, and quantified myelin sheaths at 4 dpf. Similar to results shown in Fig. 2, oligodendrocytes in *mzfmr1^-/-^* larvae expressing the control *myrf:*EGFP-CAAX reporter developed sheaths of diminished length compared to the wild-type control group (Fig. 4A,C). Expression of FMR1-IRES-EGFP-CAAX in wild-type did not impact sheath development (Fig. 4B). However, targeted FMR1-IRES-EGFP-CAAX expression in oligodendrocytes of *mzfmr1^-/-^* larvae restored individual and cumulative myelin sheath length (Fig. 4D). These results demonstrate an intrinsic oligodendrocyte-requirement for FMRP in sheath growth.

**Figure 4.**
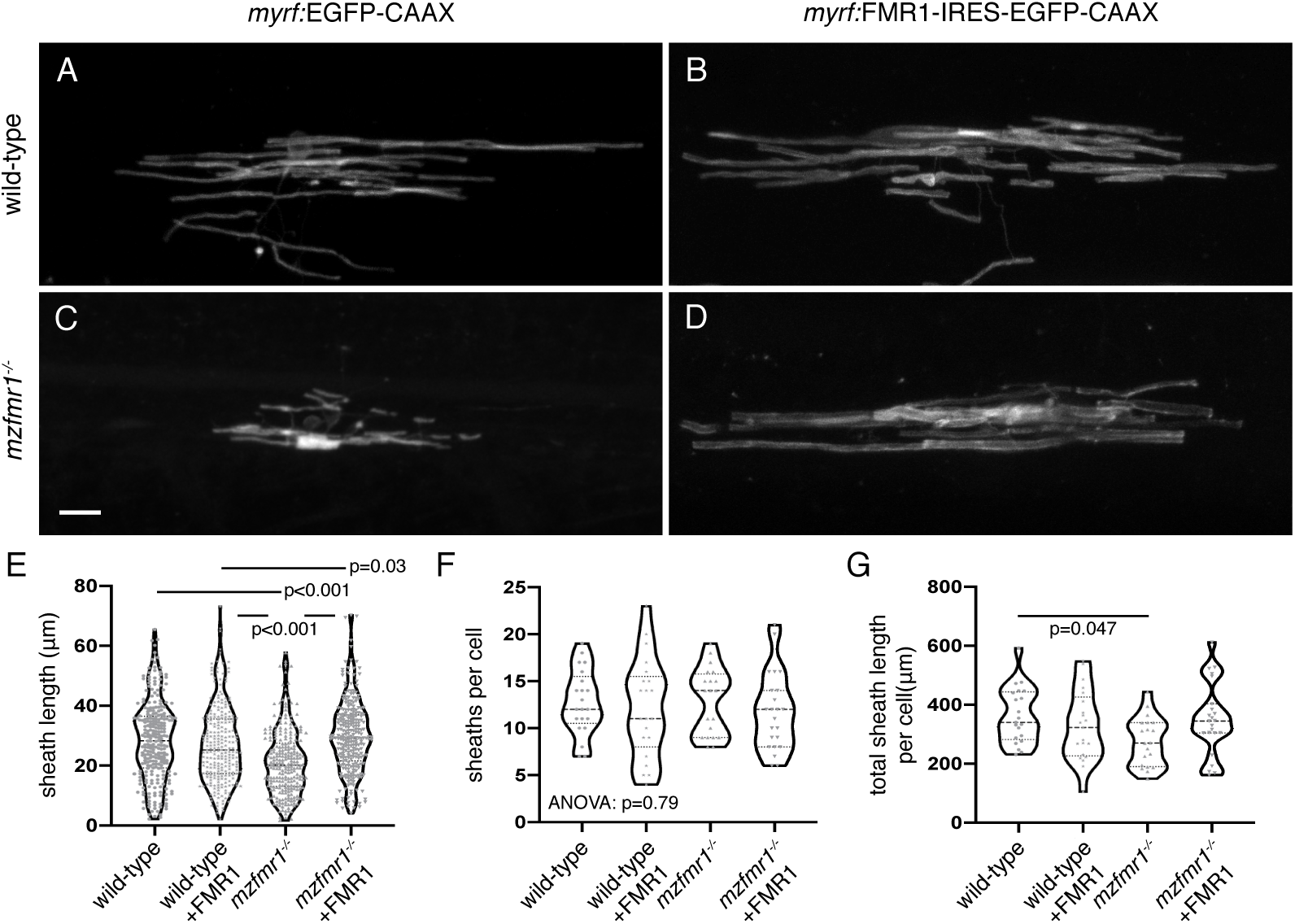
FMRP autonomously promotes myelin sheath growth. Images of living wild-type (**A**) and *mzfmr1^-/-^*loss-of-function mutant (**B**) larvae expressing *myrf:*EGFP-CAAX in oligodendrocytes. Oligodendrocyte-targeted expression of FMR1-IRES-EGFP-CAAX in wildtype (**C**) and *mzfmr1^-/-^* mutant (**D**) larvae. Average sheath length (**E**), sheaths per cell (**F**), and cumulative sheath length (**G**) in each condition. Wild-type control: 268 sheaths, 21 cells; wild-type+FMR1: 255 sheaths, 21 cells; *mzfmr1*^-/-^: 257 sheaths, 20 cells; *mzfmr1*^-/-^+FMR1: 270 sheaths, 23 cells. Significance determined by Kruskal-Wallis test with Dunn’s multiple comparisons test for average sheath length, ANOVA for sheath number, and ANOVA with Tukey’s test for and cumulative sheath length. Scale bar=10 µm.

### Loss of FMRP function leads to reduced Mbp protein expression but does not impact oligodendrocyte quantity

Oligodendrocytes in *fmr1^-/-^* mutants develop diminished myelin sheaths, which may suggest that FMRP regulates the expression of myelin proteins. We focused on Myelin basic protein (Mbp), encoded by mRNA that previous studies have shown to be a target of FMRP [9, 37]. Although FMRP can repress translation of *Mbp* mRNA in vitro [9, 37], Mbp expression was reduced in early postnatal cerebellum of *Fmr1* knockout mice [28]. To further investigate an in vivo role for FMRP, we labeled sagittal sections of transgenic larvae expressing the membrane reporter *sox10:*mRFP with antibody against Mbp, focusing on the dense myelin tracts of the hindbrain (Fig. 5A,B). Overall, we found that Mbp protein was reduced by ∼29% in *fmr1^-/-^* myelin tracts at 4 dpf (Fig. 5C). Interestingly, we also noted a regional difference in Mbp expression, in that the ventral myelin tract was most profoundly impacted by the loss of FMRP (brackets; Fig. 5D). In contrast, Mbp expression was more comparable in the dorsal myelin tract (arrowheads; Fig. 5E). These results demonstrate reduced Mbp expression in *fmr1^-/-^* embryos, which suggests a crucial role for FMRP in myelin protein production.

**Figure 5.**
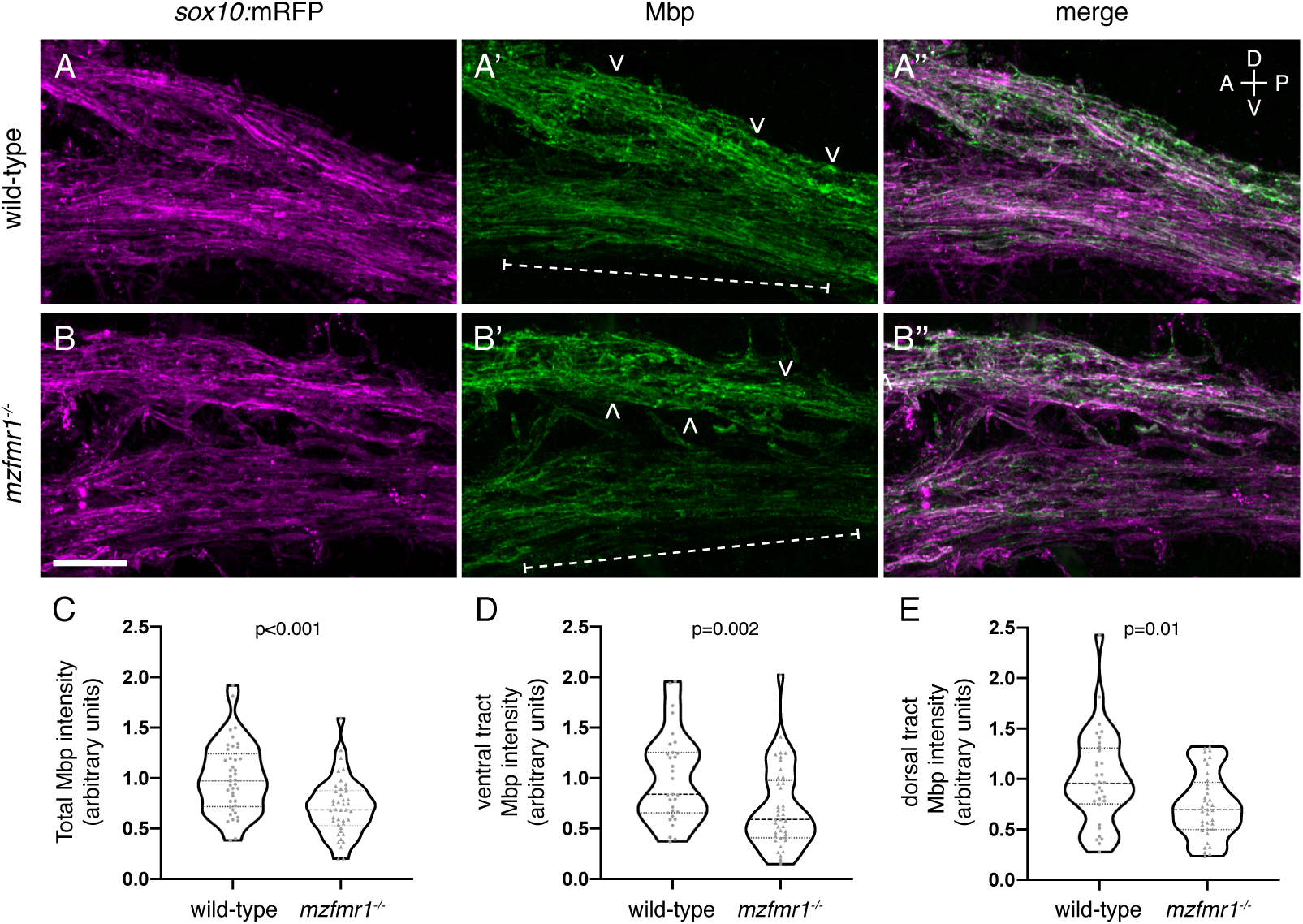
Loss of FMRP function leads to reduced Mbp protein expression. Sagittal sections through the hindbrain of *Tg(sox10:mRFP)* wild-type (**A-A’’**) and *mzfmr1^-/-^* loss-of-function mutant (**B-B’’**) larvae, labeled with antibody against Mbp. Mbp strongly co-localizes with *sox10:*mRFP+ myelin sheaths in the hindbrain. Mbp expression was quantified in both ventral (brackets) and dorsal myelin tracts (arrowheads). (**C**) Cumulative normalized quantification of Mbp intensity (wild-type n=41 sections, 30 larvae; *mzfmr1^-/-^* n*=*42 sections, 30 larvae; unpaired *t* test). (**D**) Normalized Mbp intensity in ventral hindbrain myelin tracts (wild-type n=31 sections, 26 larvae; *mzfmr1^-/-^* n*=*41 sections, 28 larvae; Mann-Whitney test). (**E**) Normalized Mbp intensity in dorsal hindbrain myelin tracts (wild-type n=32 sections, 26 larvae; *mzfmr1^-/-^* n*=*37 sections, 28 larvae; Mann-Whitney test). mRNA abundance in *fmr1^-/-^*mutants was normalized to wild-type. Scale bar=10 µm. A=anterior, P=posterior, D=dorsal, V=ventral.

Reduced Mbp abundance in *fmr1^-/-^* mutants could stem from deficient Mbp production in individual oligodendrocytes or from a reduction in the total number of cells. To investigate the latter possibility, we quantified the number of mature oligodendrocytes in the spinal cord of transgenic wild-type and *fmr1^-/-^*embryos expressing *mbpa:*EGFP-CAAX, and *sox10:*tagRFP, a cytosolic reporter of the oligodendrocyte lineage [38] (Fig. 6). We quantified the number of *mbpa*+/*sox10*+ cells in both the dorsal (Fig. 6A-B’) and ventral (Fig. 6C-D’) regions of the cord, finding comparable numbers between genotypes (Fig. 6E.F). The cumulative number of *mbpa*+/*sox10*+ cells was also unaffected by FMRP loss (Fig. 6G). These data indicate that FMRP does not regulate the quantity of oligodendrocytes in the developing spinal cord.

**Figure 6.**
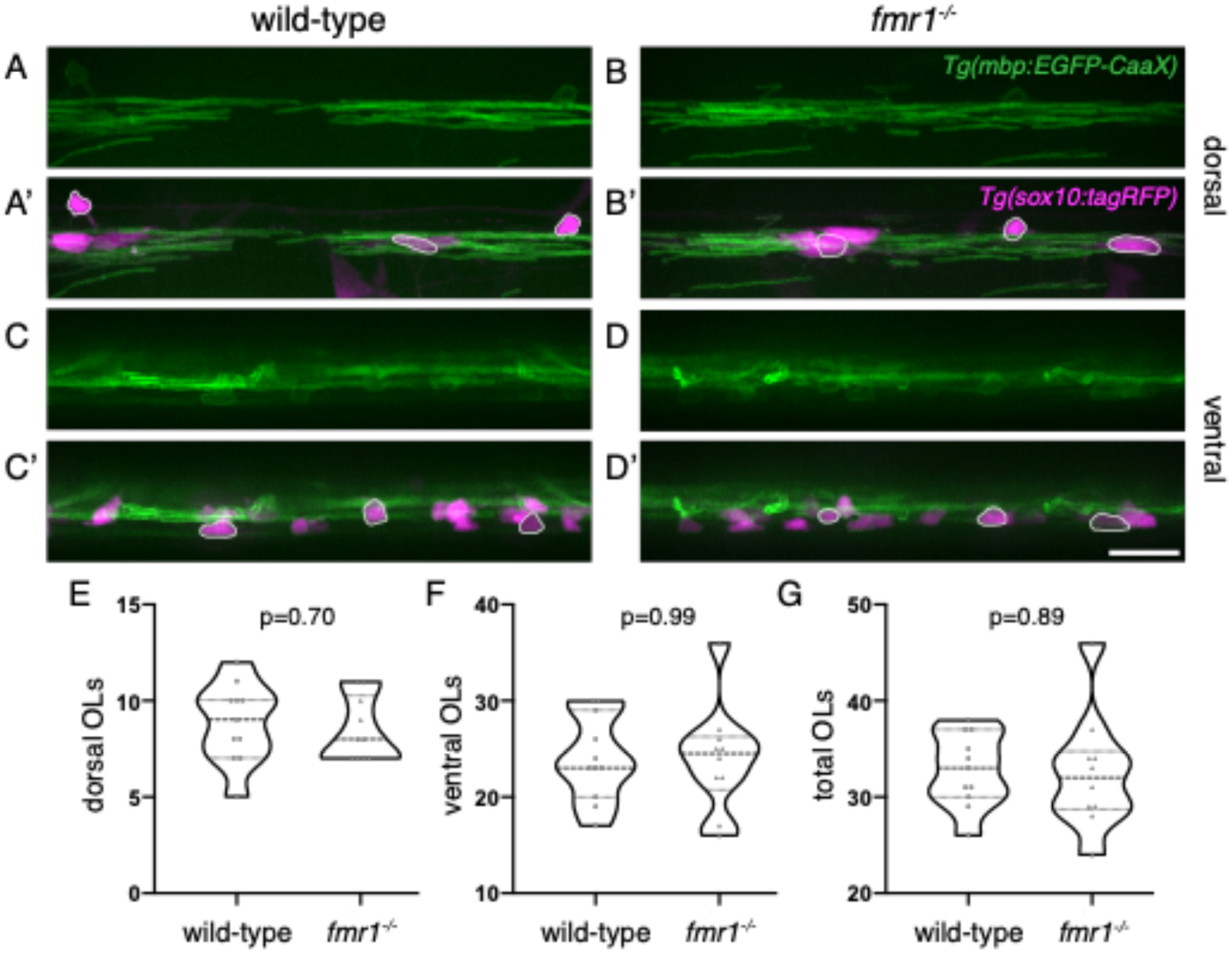
FMRP does not affect oligodendrocyte quantity in the spinal cord. Lateral images of living transgenic *Tg(mbp-EGFP-CAAX);Tg(sox10:tagRFP)* wild-type and *fmr1^-/-^* larvae at 4 dpf. *mbp+/sox10+* oligodendrocytes in the dorsal (**A-B’**) and ventral (**C-D’**) spinal cord (outlined examples in merged images). Quantification of dorsal (**E**), ventral (**F**), and cumulative oligodendrocytes (**G**)(wild-type n=11 larvae; *fmr1^-/-^*n*=*10 larvae). Statistical significance assessed using unpaired *t* tests. Scale bar=20 µm; OL=oligodendrocyte.

### Loss of FMRP does not affect mbpa mRNA abundance in myelin tracts

Because FMRP does not regulate the number of mature oligodendrocytes, reduced levels of Mbp could be due to faulty localization of *mbp* mRNA or from diminished Mbp translation. Because FMRP regulates subcellular localization of synaptic mRNAs [18], it could play a similar role for *mbp* mRNA in oligodendrocytes. To test this possibility, we performed single molecule fluorescent in situ hybridization (smFISH) [39, 40] on 4 dpf wild-type and *fmr1^-/-^ Tg(olig2:EGFP)* embryos. We quantified transcripts expressed by *mbpa*, a zebrafish ortholog of mouse *Mbp*, in individual *olig2*:EGFP+ myelin tracts in the hindbrain, which contained robust levels of *mbpa* (Fig. 7). We found that wild-type and *fmr1^-/-^* embryos contained comparable levels of *mbpa* transcript (brackets; Fig. 7C). These data indicate that FMRP does not affect *mbpa* abundance in myelin tracts, the presumptive site of Mbp translation [41].

**Figure 7.**
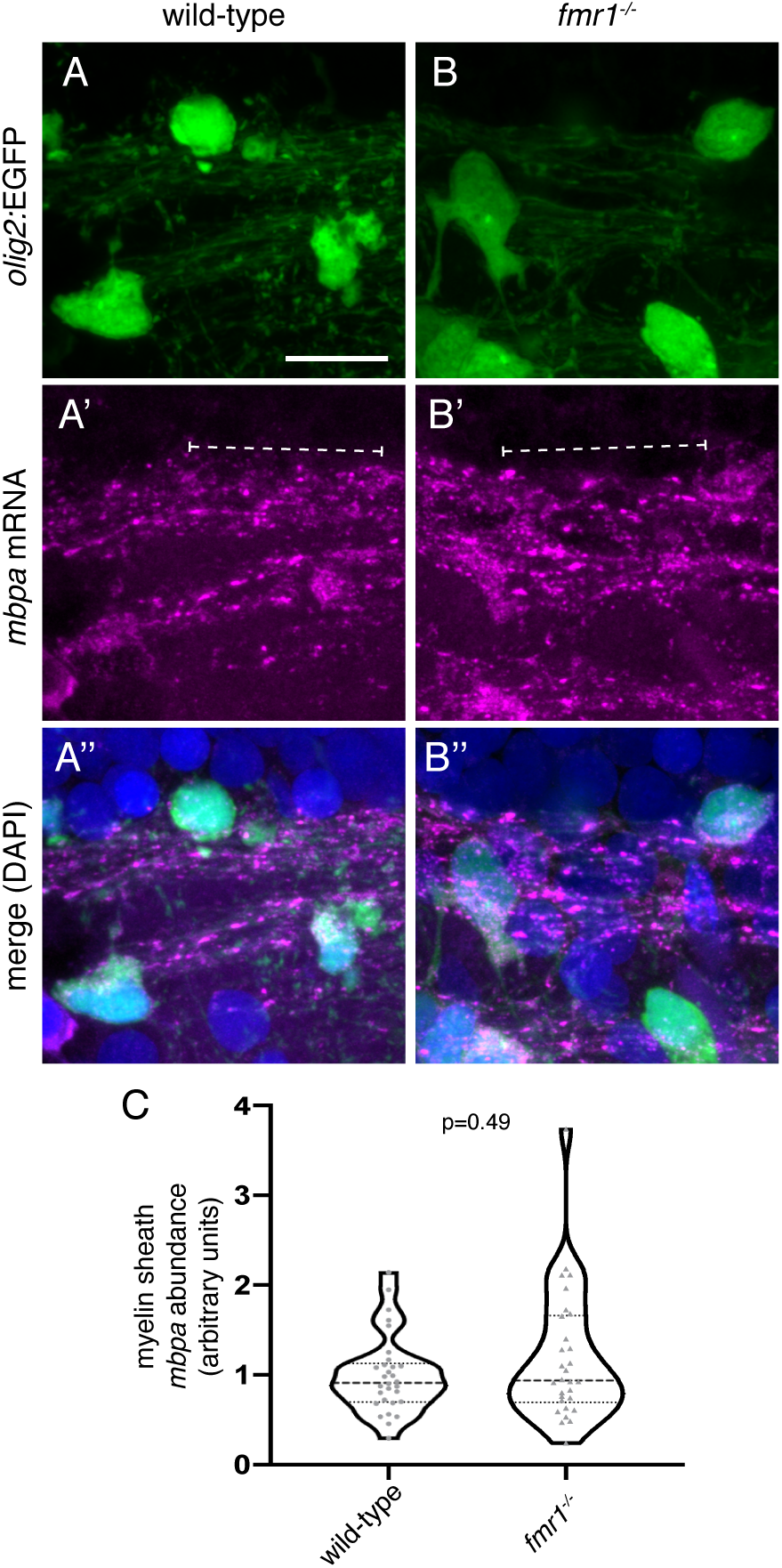
Loss of FMRP does not affect *mbpa* mRNA abundance in myelin tracts. Single molecule fluorescent in situ hybridization to detect *mbpa* mRNA in the hindbrain of *Tg(olig2:EGFP)* wild-type (**A**) and *fmr1^-/-^* (**B**) larvae. *mbpa* mRNA (**A’,B’**; brackets) is localized to *olig2:*EGFP+ myelin tracts (**A’’,B’’**). (**C**) Normalized quantification of *mbpa* mRNA abundance in hindbrain myelin tracts (wild-type n=30 myelin segments, 8 larvae; *fmr1^-/-^* n*=*30 myelin segments, 7 larvae). Statistical significance assessed with a Mann-Whitney test. mRNA abundance in *fmr1^-/-^* mutants was normalized to wild-type. Scale bar=10 µm.

### The second KH domain of FMRP is required for myelin sheath growth

FMRP contains several functional domains that mediate nuclear transport, RNA binding, and association in ribonucleoprotein (RNP) complexes (Fig. 8A), and all of these functions could contribute to myelin sheath growth. Studies of the missense mutation FMR1-I304N have demonstrated that a hydrophobic region within the second KH domain is crucial for polyribosome association [42], FMRP homodimerization, and translational regulation [43], thereby pinpointing a domain that regulates translation. We therefore tested whether this domain is required for myelin sheath growth by expressing FMR1-I304N in oligodendrocytes [44]. We injected control *myrf-*EGFP-CAAX and *myrf-*FMR1-I304N-IRES-EGFP-CAAX constructs in wild-type and *fmr1^-/-^* 1-cell embryos and examined OL morphology at 4 dpf. We found that targeted expression of FMR1-I304N in *fmr1^-/-^* mutant OLs failed to rescue the length of individual sheaths (Fig. 8E,F), but cumulative sheath length was not significantly different than wild-type (*p=*0.11; Fig. 8H). Interestingly, expression of FMR1-I304N in wild-type OLs led to a ∼30% reduction in individual myelin sheath length compared to controls (Fig. 8). Many cells in this group also developed supernumerary short branches (Fig. 8E,F), and FMR1-I304N expression in wild-type did not impact cumulative sheath length (Fig. 8G). These results demonstrate that the second KH2 domain of FMRP is essential for the growth of individual myelin sheaths but may not regulate total myelination capacity.

**Figure 8.**
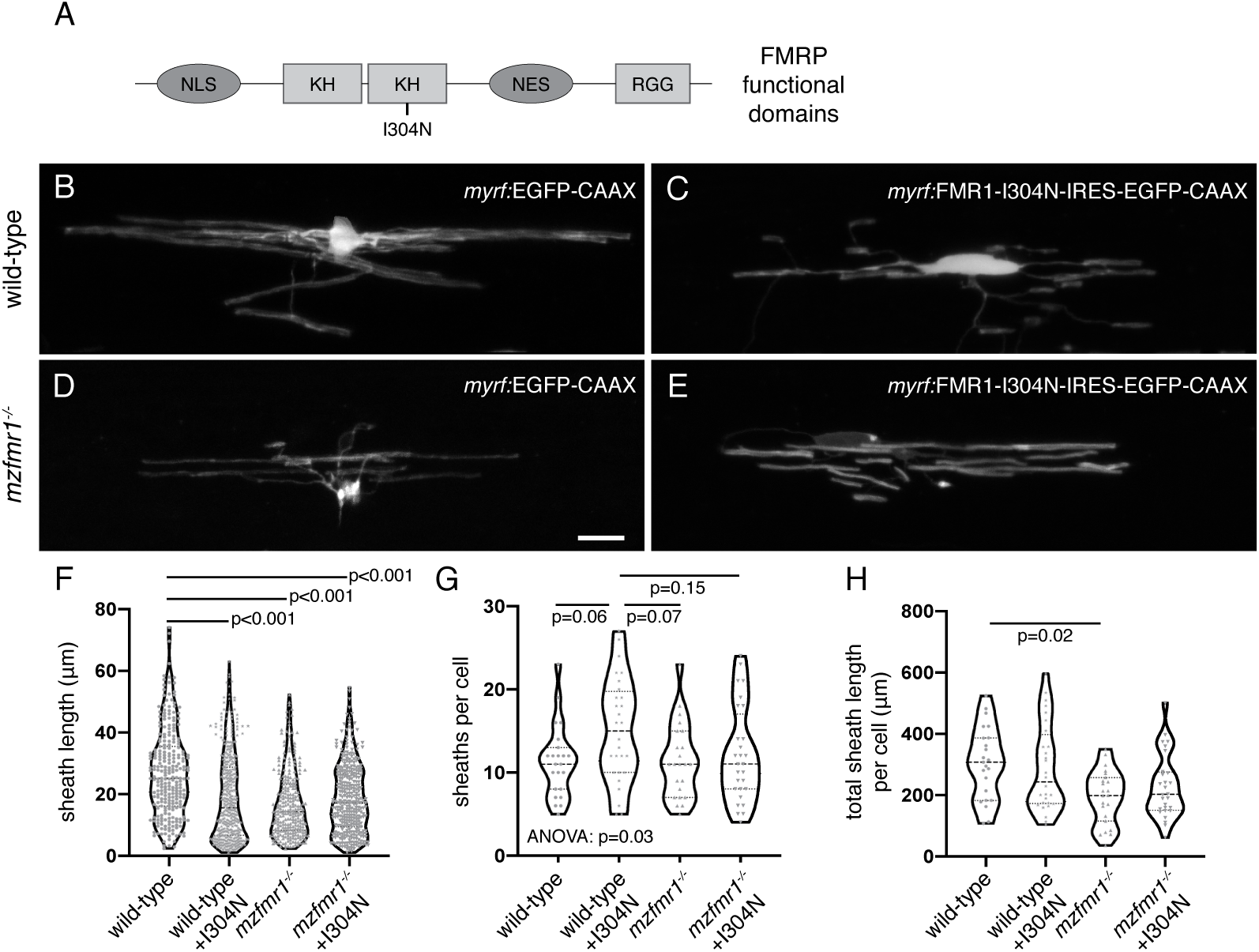
The second KH domain of FMRP is required for myelin sheath growth.(**A**) Domain structure of FMRP, including two KH regions and one RGG region (light gray boxes), as well as nuclear localization signal (NLS) and nuclear export signal (NES; dark gray boxes). The I304N missense mutation is located in the second KH domain. Lateral images of living wild-type (**B**) and *mzfmr1^-/-^* loss-of-function mutant (**C**) larvae expressing *myrf:*EGFP-CAAX in oligodendrocytes. Oligodendrocyte-targeted expression of FMR1-I304N-IRES-EGFP-CAAX in wildtype (**D**) and *mzfmr1^-/-^* mutant (**E**) larvae. Average sheath length (**F**), sheaths per cell (**G**), and cumulative sheath length (**H**) in each condition. Wild-type control: 259 sheaths, 23 cells; wild-type+FMR1: 422 sheaths, 28 cells; *mzfmr1*^-/-^: 260 sheaths, 23 cells; *mzfmr1*^-/-^+FMR1: 335 sheaths, 28 cells. Statistical significance assessed by Kruskal-Wallis test with Dunn’s multiple comparisons test for average sheath length and cumulative sheath length and ANOVA with Tukey’s test for sheath number. Scale bar=10 µm.

## Discussion

Fragile X syndrome white matter abnormalities have been attributed to primary neuronal dysmorphia or dysfunction [22]. We find that oligodendrocytes in *fmr1^-/-^* mutants develop individual myelin sheaths of reduced length as well as a dramatic decrease in total myelin. These phenotypes could result from changes in axon structure or physiology, mistargeting of oligodendrocyte processes toward axons that are not normally wrapped in myelin, or an autonomous oligodendrocyte requirement for FMRP. Although we cannot completely rule out axonal contributions, our results suggest an intrinsic oligodendrocyte requirement, because targeted expression of FMR1 can rescue myelin sheath growth in *fmr1^-/-^* mutant embryos.

FMRP can regulate several crucial steps in the lifecycle of an mRNA, including localization [18], stabilization [23], and translational control [25]. At subcellular resolution, we show that the developing hindbrain myelin tracts of both wild-type and *fmr1^-/-^* mutant embryos contained comparable levels of *mbpa* mRNA. However, Mbp protein expression was reduced in *fmr1^-/-^* mutants, which is consistent with data from the *Fmr1* knockout mouse [28]. Taken together, FMRP appears to promote Mbp translation in sheaths downstream of mRNA localization. Both in vivo studies run contradictory to prior in vitro demonstrations of translational repression of *Mbp* by FMRP [9], though it is important to note that FMRP does not always repress translation of mRNA targets: FMRP appears to promote translation of some NMDA receptor subunits, scaffolding proteins, and the immediate early gene Arc, all of which are reduced in prefrontal cortex of *Fmr1* knockout mice [45]. Intriguingly, many of these targets are also expressed in the oligodendrocyte lineage [30], and could contribute to myelin sheath growth and stabilization. FMRP plays various roles in mRNA regulation and translational control and uncovering precise in vivo requirements in developing subcellular compartments remains challenging.

Our future work will investigate the additional functional domains of FMRP to better pinpoint regions that determine localization, stabilization, and translational regulation, along with the specific target sites on mRNAs that are bound by these domains. FMRP may act to stabilize and prevent decay of *mbpa*, as detected in neuron-glia co-cultures [23]. Association of *Mbp* mRNA with an additional RBP, Quaking (Qki), appears to localize *Mbp* mRNA [46], which could explain why myelin-localized *mbpa* abundance was unaffected by the lack of FMRP. FMRP could also function more directly in translational regulation of sheath-promoting mRNAs within larger ribonucleoprotein complexes [42, 47], as introduction of FMR1-I304N fails to rescue the growth of individual sheaths in *fmr1^-/-^* mutants and induces reduced sheath growth in wild-type. Therefore, our data predict that translational control underlies the specific FMRP contribution to individual sheath growth. The precise dissection of FMRP function in mRNA regulation will require careful in vivo examinations of both FMRP binding domains and the requisite sequence motifs on mRNA targets.

Taken together, our experiments indicate a vital role for FMRP in the growth of nascent myelin sheaths. As myelination is influenced by axonal activity [48, 49], we speculate that the short myelin sheaths noted in *fmr1^-/-^* mutants are analogous to the immature dendritic spines noted in FXS patients and animal models [33, 50]; FMRP may serve as part of an activity-dependent mechanism that responds to axonal activity and initiates ensheathment. Our time-lapse experiments indicate that FMRP promotes sheath growth and dynamics during the height of axonal ensheathment by oligodendrocytes. These complex processes likely require a host of proteins, many of which may be regulated by FMRP as mRNA targets [51]. Our future work will explore additional FMRP targets driving sheath growth, including mRNA candidates that encode synaptogenic proteins. FMRP binds dozens of transcripts encoding synaptic proteins [44, 52], many of which are also expressed in the oligodendrocyte lineage [30]. Interestingly, our lab has shown that disruption of synaptic protein function in oligodendrocytes can lead to excess myelin sheaths of diminished length [35], which is comparable to FMR1-I304N overexpression (Fig. 8). Shared mechanisms of mRNA regulation in both neurons and glial would only expand the essential roles for RBPs in neural circuit development.

## Materials and Methods

### Zebrafish lines and husbandry

The Institutional Animal Care and Use Committee at the University of Colorado School of Medicine approved all animal work. Embryos were raised at 28.5°C in embryo medium and staged as hours or days post fertilization (hpf/dpf) according to morphological criteria [53]. Zebrafish lines used in this study included *fmr1^hu2787^* [34], *Tg(sox10:mRFP)^vu234^* [32], *Tg(sox10:tagRFP)^co26^* [38]*, Tg(mbp:EGFP-CAAX)^co58^*[54], and *Tg(olig2:EGFP)^vu12^* [55]. All other constructs were delivered by transient transgenesis to facilitate sparse labeling and single cell analysis.

### Plasmid construction

Multisite Gateway cloning was utilized to generate Tol2 expression plasmids, which were injected into single cell embryos along with Tol2 mRNA to produce transient transgenic animals. Human *FMR1 and FMR1-I304N* were amplified from *pFRT-TODestFLAGHAhFMRPiso1*, a gift from Thomas Tuschl [44], using the following attB primers (annealed sequence in lowercase): forward, GGGGACAAGTTTGTACAAAAAAGCAGGCTTAatggaggagctg;reverse, GGGGACCACTTTGTACAAGAAAGCTGGGTTgggtactccattcacgagtggt. We generated *pME-FMR1* and *pME-FMR1-I304N* entry clones via BP recombination with pDONR221 backbone. Middle entry clones were then cloned behind either *sox10* or *myrf* promoters, either in frame with EGFP to generate a fusion protein or with IRES-EGFP-CAAX to label cell membranes with a distinct peptide.

Entry clones were LR-recombined with either the *pDEST-Tol2-CG2* destination vector (green heart marker; for *pEXPR-myrf:FMR1-IRES-EGFP-CAAX* and *pEXPR-myrf:FMR1-I304N-IRES-EGFP-CAAX*) or *pDEST-Tol2-pA2* vector for *pEXPR-sox10:FMR1-EGFP*.

Published plasmids: *p3E-7.2sox10* [31], p5E*-myrf*, *p3E-IRES-EGFP-CAAX*, *pEXPR-tol2-mbp:EGFP-CAAX* [35]*, p3E-EGFP* [56].

### Imaging and analysis

With the exception of smFISH and IHC, live larvae were imaged in all experiments. Larvae were embedded laterally in 1.2% low-melt agarose containing 0.4% tricaine for immobilization. We acquired images on a Zeiss CellObserver SD 25 spinning disk confocal system for time-lapse microscopy and cell counts (Carl Zeiss) or a Zeiss LSM 880 for all other experiments (Carl Zeiss). Images were captured with Zen software (Carl Zeiss), then processed and analyzed using Fiji/ImageJ or Zen Blue (Carl Zeiss).

### smFISH probe design

*mbpa* smFISH probes were designed using the Stellaris RNA FISH Probe Designer tool by entering the zebrafish *mbpa* cDNA sequences obtained from Ensemble Genome Browser from transcript *mbpa-206* (GRCz11). Probes with highly repetitive sequences were removed. The probes were ordered with a CAL Fluor® Red 610 Dye. Probes were resuspended in Tris-EDTA, pH 8.0 and stored at a stock concentration of 12.5 µM at −20°C.

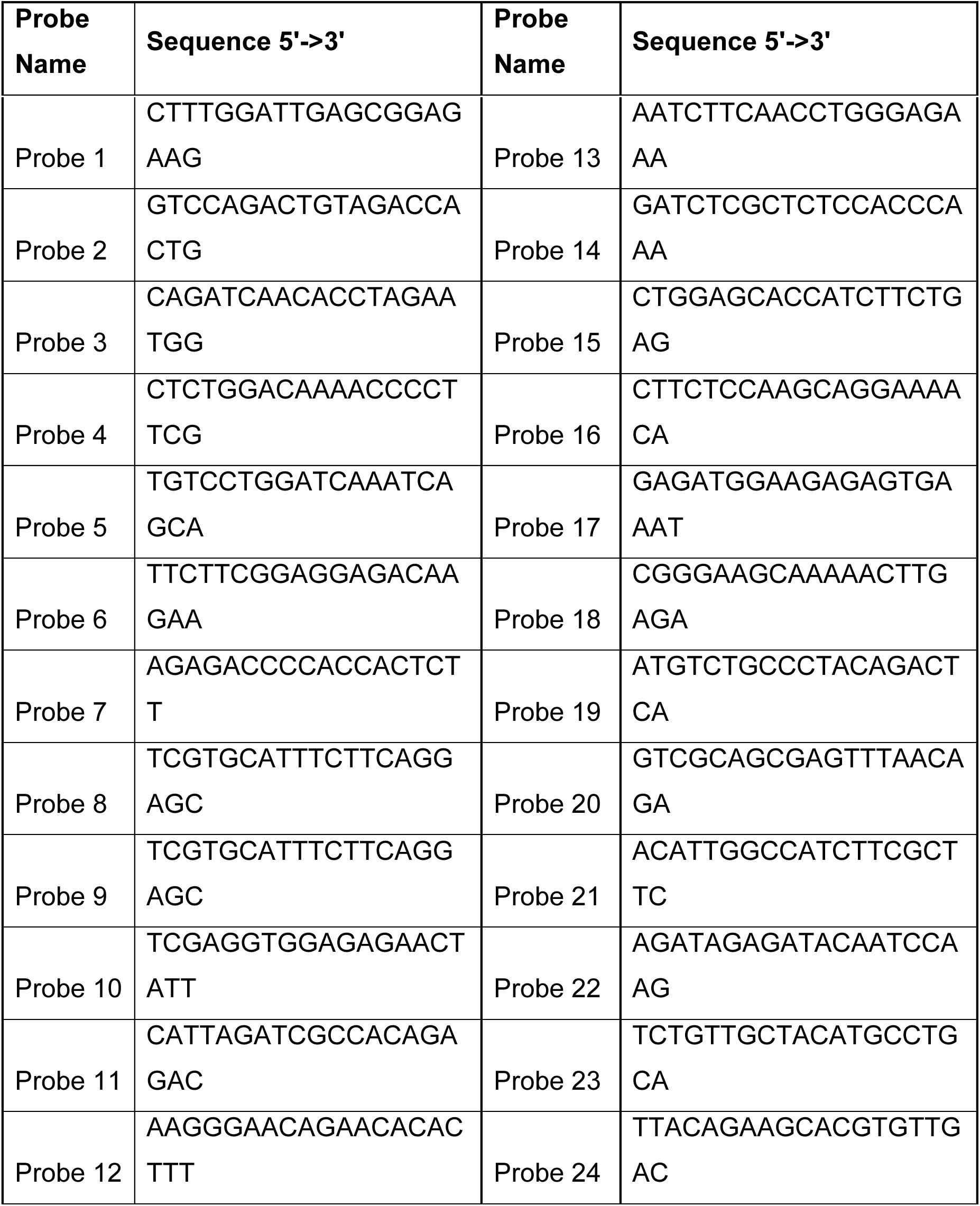

### smFISH experimental procedure

The smFISH protocol was adapted from three published protocols [57–59]. First, larvae were sorted for *olig2:*EGFP expression and fixed O/N in 4% paraformaldehyde at 4°C. Larvae were embedded laterally in 1.5% agar, 5% sucrose blocks and transferred to a 30% sucrose solution O/N at 4°C. Blocks were frozen on dry ice and sectioned with a Leica cryostat into 20 µm thick sections and placed on microscope slides (Fisherbrand cat #: 12-550-15). Slides were not allowed to dry more than 5 min before adding 4% paraformaldehyde to fix the tissue at room temperature (RT) for 10-20 min. The slides were quickly rinsed with 1X PBS twice. The tissue was permeabilized with 70% cold ethanol at −20°C for 2 hours. Parafilm was placed over tissue to prevent evaporation at all incubation steps. The tissue was rehydrated with wash buffer (10% DI formamide, 2XSSC in molecular grade water, 1 mL DI formamide, 1 mL 20X SSC) for 5 min at RT. From this point on, care was taken to protect the tissue and probes from light as much as possible. Hybridization Buffer was made: 2XSSC, 10% DI formamide, 25mg/mL tRNA, 50mg/mL bovine serum albumin, 200mM ribonucleoside vanadyl complex in DEPC water. Aliquots were made and stored at −20°C. Probe was added to hybridization buffer 1:100 for a final concentration of 125 nM. As a control, slides were incubated in hybridization buffer with no probe. Slides were quickly rinsed with fresh wash buffer followed by 2 wash steps at 37°C for 30 minutes. DAPI was added at 1:1000 concentration in wash buffer to the tissue for 7 min at RT. Slides were quickly rinsed twice with wash buffer. Finally, slides were mounted with Vectashield mounting media and a No. 1 coverslip and sealed with nail polish. All slides were stored and protected from light at 4°C.

### smFISH microscopy

Images of smFISH experiments were obtained using a Zeiss LSM 880 with Airyscan confocal microscope and a Plan-Apochromat 63x, 1.4 NA oil immersion objective. The acquisition light path used Diode 405, Argon 488, HeNe 594 lasers, 405 beam splitter and 488/594 beam splitters, and Airyscan super resolution detector. Imaging was performed using Zeiss Zen Black software and parameters included: 1024×1024 frame size, 1.03 µsec pixel dwell time, 16-bit depth, 2.4x zoom. 1.8% 488 laser power, 5% 594 laser power, 0.5% 405 laser power, 705 gain, and z intervals of 0.3 µm. All images were taken in the hindbrain of zebrafish larvae. Myelin segments were selected for imaging based on expression of *olig2:*EGFP and Quasar-610 fluorescence. Post-image processing was performed using Airyscan Processing set to 6.8.

### Immunohistochemistry

4 dpf *Tg(sox10:mRFP)* embryos were fixed in 4% paraformaldehyde/1xPBS overnight at 4°C. Embryos were washed 3×5 minutes in 0.1%Tween/1xPBS (PBSTw), rocking at room temperature (RT). For antigen retrieval, embryos were placed in 150mM Tris pH 9 for 5 minutes at RT and then 15 minutes at 75°C. Embryos were rinsed in 1xPBS/0.1%Tween, then embedded in 1.5% agar/30% sucrose and immersed in 30% sucrose overnight. Blocks were frozen on dry ice and 20µm sagittal sections were taken with a cryostat microtome and collected on polarized slides. Slides were mounted in Sequenza racks (Thermo Scientific), washed 3×5 minutes in 0.1%Triton-X 100/1xPBS (PBSTx), blocked 1 hour in 2% goat serum/2% bovine serum albumin/PBSTx and then placed in primary antibody (in block) overnight: rabbit α-Mbp (1:200; [60]). Sections were washed 1.5 hours in PBSTx, and then incubated 2 hours at RT in secondary antibodies: AlexaFluor 488 goat α-rabbit. Sections were washed for 1 hour in PBSTx, incubated with DAPI (1:2000 in PBSTx) for 5 minutes, washed 3×5 minutes in PBSTx, then mounted in Vectashield (Vector Laboratories).

### Immunohistochemistry microscopy

Images of immunohistochemistry experiments were obtained with the same settings as smFISH, along with the following changes: 512×512 frame size, 0.85 µsec pixel dwell time, 16-bit depth, 1.8x zoom. Line averaging was set to 2, 6.5% 488 laser power, 4.5-5.5% 594 laser power, 1% 405 laser power, 619 gain, and z intervals of 0.144 µm. All images of single cells were taken in the hindbrain of zebrafish larvae. Cells were selected for imaging based on *sox10:*mRFP fluorescence. Post-image processing was performed using Airyscan Processing set to 1.0.

### Quantification and Statistical Analysis

#### smFISH quantification

All quantification was performed in ImageJ Fiji using a custom script created by Karlie Fedder (available upon request). First, z intervals were selected for myelin tracts using the “Make Substack” feature in Fiji. Substacks of myelin tracts in the hindbrain included 13 steps with an interval of 0.3 µm. Each substack was maximum z-projected. Background was subtracted using a 2.5 rolling ball. The image was then thresholded by taking 3 standard deviations above the mean fluorescence intensity. Puncta were analyzed using the “Analyze Particles” feature with a size of 0.01-Infinity and circularity of 0.00-1.00. Using the maximum projection of the *olig2:*EGFP channel, a region of interest (ROI) was drawn around myelin segments using the rectangle tool to draw a square of 100×100 pixels (4.39um x 4.39um). All thresholded puncta were inspected to ensure single molecules were selected. Occasionally, threshold puncta fell on the border of the ROI and these were excluded from measurements. *mbpa* transcripts are highly expressed and counting individual puncta was not consistently reliable. Therefore, to measure each puncta, we overlaid the thresholded image on the maximum z projected image and calculated the integrated density (area x average fluorescence intensity) using the “IntDen” measurement.

### Immunohistochemistry quantification

First, z intervals were selected for hindbrain myelin tracts using the “Z Project” feature in Fiji. Substacks of myelin tracts in the hindbrain included 20 steps with an interval of 0.144 µm. Each substack was projected via “Sum Slices”. Fluorescence channels were split and saved as 16-bit images. Next, the mRFP channel was duplicated and automatically thresholded using the Otsu method. The wand tracing tool was used to select myelin tracts and intensity measurements were taken by redirecting to the Mbp channel (Analyze, Set Measurements, Redirect). Following quantification of the entire dataset, individual values were normalized to average wild-type Mbp expression levels.

### Statistics

All statistics were performed in Graphpad Prism (version 8). Outliers were identified using a ROUT with Q=1%. Normality was assessed with a D’Agostino and Pearson omnibus test. For two groups, unpaired comparisons were made using either unpaired two-tailed *t* tests (for normal distributions) or Mann-Whitney tests (abnormal distributions). For multiple comparisons, an ANOVA was followed with Tukey’s individual comparisons to each mean for normal distributions or Kruskal-Wallis test was followed with Dunn’s individual comparisons to each mean for abnormal distributions.

## Acknowledgments

The authors would like to thank Matthew Taliaferro for insightful conversation. This work was supported by US National Institute of Health (NIH) grant R01 NS095679 and a gift from the Gates Frontiers Fund to B.A. and NIH grant R21 NS110213 to C.D. and B.A.

## Author Contributions

C.D. and B.A. conceived the project. K.Y. and C.D. performed FISH experiments. C.D. performed all additional experiments and collected all data. K.Y. analyzed FISH data, C.D. analyzed all other data. C.D. wrote and B.A. edited the manuscript.

